# Epigenome-wide analysis reveals novel DNA methylation signatures significantly associated with the infant pupillary light reflex, a candidate intermediate phenotype for autism

**DOI:** 10.1101/2025.01.10.632385

**Authors:** Laurel A. Fish, Teodora Gliga, Anna Gui, Jannath Begum Ali, Luke Mason, Mark H. Johnson, Tony Charman, Terje Falck-Ytter, Emily J. H. Jones, Radhika Kandaswamy, Francesca Happé, Chloe C. Y. Wong

## Abstract

Autism is a highly heterogeneous neurodevelopmental condition, currently diagnosed based on behavioural characteristics. Candidate early intermediate phenotypes, such as the Pupillary Light Reflex (PLR), a reflexive constriction of the pupil in response to increased optical luminance, may provide insights into etiological factors and potential biomarkers, such as DNA methylation (DNAm), involved in the emergence of autism. We conducted epigenome-wide DNAm association analyses of 9-, 14-, 24-month PLR onset latency and constriction amplitude in a sample of 51 infants enriched for autism family history, using buccal DNA collected at 9-months.

Our epigenome-wide analysis (EWAS) identified four stringently significant differentially methylated probes (*p* < 2.4 × 10^−7^) associated with cross-section PLR latency measurements at 14– and 24-months, and with 14-to 24-month PLR latency developmental change. Differentially methylated probes associated with PLR amplitude were identified, but at a less stringent discovery threshold (*p* < 5 × 10^−5^). Our region analyses identified several significant differentially methylated regions associated with both PLR latency and amplitude Downstream exploratory pathway analysis identified enrichment for multiple developmental biological processes, as well as several susceptibility genes to autism and related neurodevelopmental conditions including *NR4A2, HNRNPU and NAV2*. Our findings provide novel insight into the role of DNAm in PLR development and illuminate biological mechanisms underpinning altered PLR in infancy in emerging autism.

---

Autism is a life-long highly heterogenous neurodevelopmental condition characterised by difficulties in social interactions and communication, restricted/repetitive interests/behaviours and sensory anomalies (1). Autism cannot be diagnosed reliably until two years, at the earliest, when behavioural manifestations are sufficiently developed for clinical observation (2). The aetiology of autism is not fully understood, but prospectively investigating early intermediate phenotypes, defined as pre-diagnosis measurable features associated with the later behavioural characteristics of the condition (3), could be fruitful in understanding early autism development (4). The pupillary light reflex (PLR) is a candidate early intermediate phenotype for autism (5). The PLR is the reflexual constriction of the pupil in response to increased optical luminance (6) and is primarily mediated by a relatively simple four-neuron visual sensorimotor circuity (7). This well-studied index of autonomic functioning is governed by the parasympathetic and sympathetic pathways (8) and its regulation involves neural signal transduction and cholinergic and norepinephrine neurotransmission (8–12). Recent findings provide strong evidence to suggest the implication of the PLR, thus the neural underpinnings of this reflex, in the emergence of autism, indicating the PLR as a candidate biomarker for autism. Cross-sectionally, 10-month-old infants with a family history of autism have a larger and faster PLR relative to controls (13), with larger 10-month PLR constriction amplitude associating with later diagnosis (14). Developmentally, from 6 to 24 months, PLR tends to become faster and larger (15,16), but a decreasing pattern of pupil constriction amplitude over time in those who later receive an autism diagnosis has been demonstrated (14). This finding was replicated in a partially overlapping sample in which a decrease in amplitude between 9– and 14-month was associated with increased autism-associated traits (5). In the same sample, those with a later diagnosis demonstrated a significantly steeper decrease in latency from 14 to 24 months relative to those with other outcomes (5). Additionally, infants with a higher polygenic score for autism was associated with a smaller decrease in latency in the first year (5).

Increasing evidence suggests autism arises during early development from complex interactions between genetic and environmental factors (17). Though the path through which these factors coalesce remains unclear, a growing body of evidence points to altered epigenetic signatures in autism (18). Epigenetic modifications, such as DNA methylation (DNAm), are chemical markers that can influence gene expression without altering the underlying DNA sequence itself (19). DNAm is a key biological mechanism involved in the timed up-or down-regulation of gene expression (20,21). Recent epigenome-wide association studies (EWAS) using both post-mortem brain tissue and peripheral tissue (e.g. blood, buccal or saliva) from living participants have identified several differentially methylated sites and regions, as well as co-methylated modules, associated with both autism diagnosis (22–24) and dimensional autism traits (25). Autism-associated DNAm signatures have been identified in genes and pathways linked to critical functional processes, including neurodevelopment, immune response, synaptic functioning, and microscopic neural structural variations, highlighting their potential and relevance as valuable biomarkers for autism (22–24,26,27).

Notably, all the above investigations used samples taken post-diagnosis. Autism is a neurodevelopmental condition, thus prospectively investigating DNAm before diagnosis may be more fruitful in mapping the emergence of the disorder. A recent EWAS (28) identified differential peripheral tissue DNAm in 8-month-olds associated with 3-year autism diagnosis and dimensional autism traits. However, these associations were not statistically significant after stringent epigenome-wide multiple testing corrections. An alternative, potentially more powerful, method of exploring the role of DNAm in the emergence of autism is to investigate the association of DNAm with early intermediate phenotypes (4), which tend to be placed at an earlier stage in the aetiological pathway (3). Indeed, EWAS with two candidate 8-month intermediate phenotypes of social attention (face looking and neural response to faces) was recently conducted, though, again, no probe-wise associations were significant after epigenome-wide multiple testing corrections (28). Given PLR is a quantifiable reflex primarily involving a much simpler neuro-circuitry, it is likely placed at an earlier stage in the aetiological pathway for autism than biomeasures of social attention. This makes it an ideal alternative candidate intermediate phenotype to explore the role of DNAm in the emergence of autism.

In this study, we present findings from epigenome-wide DNAm association analysis of infant PLR onset latency and constriction amplitude, using buccal tissue taken from 51 9-month-old infants enriched for autism family history. We applied an epigenome-wide approach, to explore the association of probe– and region-level DNAm with variability in cross-sectional (9-, 14-, and 24-month) and developmental (9-to 14-month and 14-to 24-month changes) PLR onset latency and constriction amplitude. Using the identified probes and regions, we conducted gene ontology enrichment analysis to uncover biological significance and performed functional exploratory analyses to investigate genetic and epigenetic links to autism.

## Methodology

### Sample

Participants in the current analysis were recruited into the British Autism Study of Infant Siblings (BASIS; www.basisnetwork.org; SM 1), a longitudinal study of the development of infants at an increased familial likelihood for receiving a later diagnosis of autism due to having a first degree relative with a diagnosis of autism.

The current sample consisted of a total 51 male infants who had DNA methylation data quantified at 9-months, and PLR data from at least one of the following time points: 9-months, 14-months or 24-months. The sample was enriched for infants with an elevated familial likelihood for receiving a later diagnosis of autism (N=42); 9 infants had a low familial likelihood for autism having no family history of the condition. We focused specifically on this age range of PLR measurements because *1)* previous investigations have reported association of the PLR change across this period with a later autism diagnosis, and *2)* this is the key early developmental timeframe in which behavioural autism characteristics gradually emerge (29). As DNAm is dynamic over development (18), we focused on DNAm at our first PLR timepoint (9-months) to investigate the developmental sequelae in PLR following earlier DNAm and to narrow the age range of DNAm as recommended by previous investigations (30). We additionally included only male infants to minimise epigenetic heterogeneity.

### Pupillary Light Reflex (PLR)

The PLR was induced while infants passively watched stimuli that transitioned from a black to white screen on a monitor (31), see Figure 1. Pupil diameter data was collected at 9-, 14– and 24-months using Tobii T120 eye tracker (sampling rate = 60Hz; Tobii Technology AB, Danderyd, Sweden; millimetres to 2 decimal places). Pre-processing of raw pupil data (see SM 2.1) removed artefacts and extracted PLR latency (constriction onset time; ms) and amplitude (constriction magnitude relative to average pupil size before latency; %). Latency was defined as the minimum acceleration from 110-570ms after white slide onset. Amplitude was calculated as the maximum constriction within the interval 170– 1450ms post-latency, relative to the average pupil diameter size at latency onset (32) The parameter identification time windows were determined by a series of optimisations investigations. Three or more valid trials per time point were required for inclusion. The median latency and amplitude per individual per time point were calculated.

**Figure 1.**
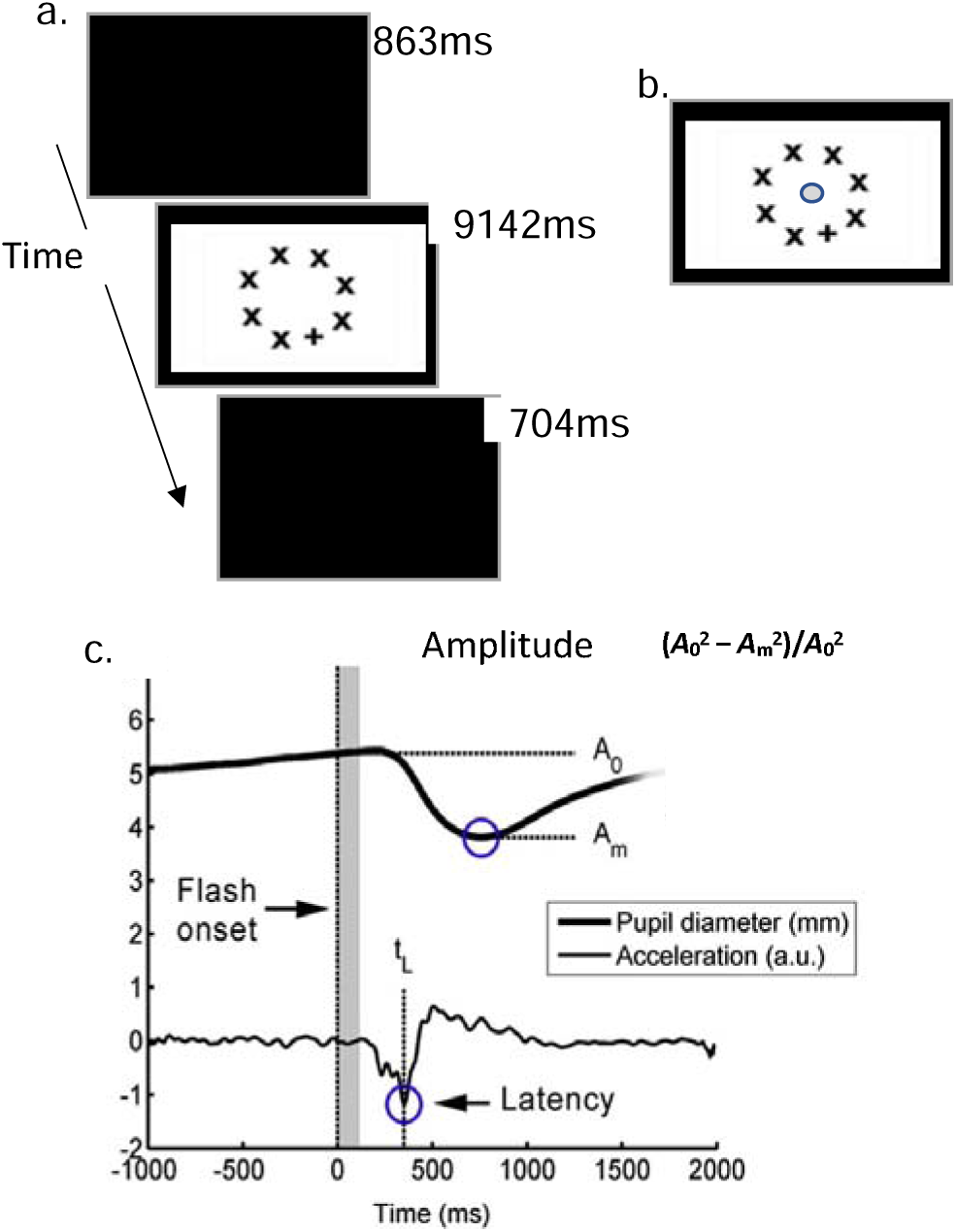
PLR stimulus and pupil response. a) Stimulus presented to the cohort, 32 times at 9/14 months and 16 times at 24 months; b) PLR inducing stimulus slide with blue circle demonstrating the approximate location of the centre area of interest (centre ∼5% of the white slide) for first look for trial to be included in the analysis; c) pupil size (mm) and acceleration (a.u.) traces demonstrating relative constriction amplitude formula components (33)*(A*_0_ = average pupil diameter in the window 100ms before constriction onset latency; *A*_m_ = minimum pupil diameter in 170-1450ms relative to *A_0_*. ms = milliseconds).

To account for the variance in the PLR parameters at each timepoint explained by the covariates of missingness (i.e., percentage of missing trials from the stimuli) and baseline pupil diameter (i.e., the average pupil diameter up to 100ms before PLR latency), we conducted linear multiple regression analysis with latency or amplitude as the dependent variables and missingness and baseline as the predictor variables (SM 2.2). Linear regressions were conducted within timepoint, and residuals from these analyses were extracted and used in all subsequent analysis as the PLR latency and amplitude variables.

In total 10 PLR phenotypes were analysed in this study which included 6 cross-sectional PLR measures (latency and amplitude at 9-, 14– and 24-month) and 4 PLR developmental change scores (9-to 14-month and 14-to 24-month for latency and amplitude). Change scores were calculated using subtraction of the older minus the younger measure. The inclusion of all 10 PLR phenotypes, encompassing both cross-sectional and developmental measures, is based on the current literature, which suggests that both latency and amplitude at various time points across infancy may be associated with autism either directly such as 10-month amplitude and latency (13,33), or as part of an associated developmental trajectory (5,13). The inclusion of these phenotypes enable a comprehensive exploration of the role of epigenetics in early PLR as an intermediate phenotype for emerging autism.

### DNA methylation (DNAm)

Buccal samples were collected at 9 months from 51 infants and processed using a standard pipeline (28). Genomic DNA (500ng) from each sample was extracted and treated with sodium bisulfite using the Zymo EZ DNA Methylation-Lightning Kit™ (Zymo Research, Irvine, CA, USA). The Illumina Infinium® HumanMethylation450 BeadChip kit (450K array; Illumina, San Diego, CA, USA) was used to assess DNA methylation at 482,421 sites (#x25A1;2 % of all CpG loci) throughout the genome. We then quantified DNAm using the HiScan System (Illumina, San Diego, CA, USA) and extracted signal intensities using Illumina GenomeStudio software (Illumina, San Diego, CA, USA) for each probe.

All epigenomic data processing and downstream analysis was conducted using R version 4.0.2. and R studio (34,35) on the high-performance computing facilities at King’s College London (CREATE). Methylation signal intensity data was imported using *methylumi R-*package (36). Data quality control and pre-processing was conducted using *wateRmelon* R-package (37). The ‘pfilter’ function removed failed samples and probes (e.g., low bead counts (< 3) or detection p-values > 0.05 for more than 1% of probes). The ‘dasen’ function was then used to normalise the data using a standardised pipeline (37) to address background correction, dye bias, probe adjustments, batch effects, and other non-biological variations related to external parameters (e.g., array order) (38). Cross-reactive and polymorphic probes, as identified in Illumina annotation files and recent studies (39,40), were excluded. The final dataset included 402,971 probes with methylation levels expressed as beta (β) values ranging from 0 (unmethylated) to 1 (fully methylated), as shown in SM 3 Figure 1.

### Statistical analysis

#### EWAS analysis

In total, 10 separate EWAS were conducted using linear multiple regressions fitted individually for each CpG. Models were built with processed DNAm β as the dependent variable and the PLR phenotype as the predictor with age, batch, 10 principal components (identified using location-based principal component analysis (41), SM 3.2) and estimated cell type proportion for 7 common cell-types in buccal samples (identified using EpiDISH R package (42), SM 3.3) as covariates. To account for account for multiple testing in epigenome-wide association studies, we applied a ‘stringent’ 450k *significance* threshold of *p* < 2.4 × 10^−7^ (43), and a more liberal ‘discovery’ *significance* threshold of *p* < 5 × 10^−5^, as commonly utilised in EWAS of complex phenotype including autisms (24,26–28,44).

#### Differential methylated region (DMR) analysis

We used *dmrff* R-package (45) to identify DMRs associated with the 10 phenotypes of interest. DMRs were determined using EWAS summary statistics and defined as regions of two or more neighbouring probes within 500 base pairs with EWAS *p-value* < 0.05 and effect estimates in the same direction (negative or positive). The p-value threshold for significant DMR was set at 0.05 after Bonferroni adjustments.

#### Downstream exploratory analysis

To assess the biological significance of genes linked to probes associated with the PLR phenotypes, we performed Gene Ontology (GO) enrichment analysis using the top 1,000 probes (ranked by p-value) and significantly associated DMRs (SM 5 Table 1). Analysis focused on biological processes and was conducted with the PANTHER overrepresentation test (46–48), referencing genes annotated to the 402,971 probes used in EWAS and DMR analyses. Gene annotations were based on Illumina UCSC hg19 annotations (49). Significant enrichment required at least two genes per term and an FDR-adjusted p-value ≤ 0.05. We investigated whether probes significantly associated with PLR in the EWAS or DMR analysis were previously linked to autism in the MRC-IEU EWAS catalogue (50) (October 2024) or located within autism-associated genes listed in the SFARI Gene database (51) (September 2024; gene.sfari.org).

**Table 1.** Summary of probe-wise EWAS and DMRs identified for each phenotype.

| Phenotype | N.<br>participants | $\lambda$ | PLR-associated DMP analysis | | PLR-associated<br>DMR analysis |
| --- | --- | --- | --- | --- | --- |
| | | | N. stringent-<br>sig. probes<br>($p<2.4 \times 10^{-7}$) | N. discovery-<br>sig. probes<br>($p<5 \times 10^{-5}$) | N. Bonf.-adj.<br>sig. DMR<br>($p>0.05$) |
| **<u>Latency (ms)</u>** |  |  |  |  |  |
| 9 months | 48 | 1.16 | 0 | 28 | 1 |
| 14 months | 47 | 1.14 | 2 | 27 | 1 |
| 24 months | 40 | 1.02 | 1 | 21 | 1 |
| 9 to 14 months | 44 | 0.94 | 0 | 6 | 1 |
| 14 to 24 months | 37 | 1.20 | 1 | 60 | 9 |
| **<u>Amplitude (%)</u>** |  |  |  |  |  |
| 9 months | 46 | 1.07 | 0 | 11 | 1 |
| 14 months | 47 | 1.03 | 0 | 21 | 1 |
| 24 months | 37 | 1.05 | 0 | 28 | 5 |
| 9 to 14 months | 42 | 1.06 | 0 | 28 | 3 |
| 14 to 24 months | 34 | 0.96 | 0 | 15 | 7 |

*Note: $\lambda$ = genomic inflation factor; sig. = significant. Bonf.-adj = Bonferroni-adjusted.*

## Results

### DNAm Analysis

Our study identified multiple differentially methylated probes and regions associated with PLR latency and amplitude. A summary of significant results from the epigenome-wide association studies (EWAS) and differentially methylated region (DMR) analyses are presented in Table 1 (SM 4.1 includes Q-Q plots and EWAS Manhattan plots).

We identified four differentially methylated probes significantly associated with three PLR latency phenotypes at a stringent p-value threshold (*p < 2.4×10^−7^*), Table 2. SM 4 Table 1 lists all probes associated with PLR latency phenotypes above the discovery p-value threshold (*p < 5 × 10^−5^*).

**Table 2.** Differentially methylated probes associated (*p < 2.4* x *10^−7^*) with PLR latency.

| Probe | Latency phenotype (months) | Genomic locations (hg19) | Illumina gene annotation | Effect size ($\beta$) | p-value |
| --- | --- | --- | --- | --- | --- |
| cg05148717 | 14 | chr6:28829171 | | -0.00055 | $3.74 \times 10^{-8}$ |
| cg22367466 | 14 | chr12:6840532 | *COPS7A* | -0.00035 | $4.28 \times 10^{-8}$ |
| cg09732535 | 24 | chr16:89331997 | | 0.00066 | $1.85 \times 10^{-8}$ |
| cg15130433 | 14 to 24 | chr6:159240081 | *EZR* | 0.00025 | $2.21 \times 10^{-7}$ |

*Note: A negative effect size (-) indicates hypermethylation associated with faster latency (for cross-sectional measurements) or latency becoming slower overtime (for change scores).*

In total, 13 DMRs significantly associated with PLR latency; each latency phonotype had a single significantly associated DMR, except 14-to 24-months PLR latency change where nine significantly associated DMRs were found. The DMRs significantly associated to each PLR latency phenotype are listed in Table 3 and SM 4 Table 2.

**Table 3.** Summary of the significantly associated DMRs for each PLR latency phenotype.

| Latency phenotype (months) | Genomic locations (hg19) | N probes | Probes | Illumina gene annotations | Effect size ($\beta$) | Bonf.-adjust p-value |
| --- | --- | --- | --- | --- | --- | --- |
| 9 | chr10:74927623-74927863 | 8 | cg15213114; cg04167018; cg21416602; cg25138168; cg08571229; cg04749667; cg16124546; cg12276298 | *FAM149B1*; *ECD* | -0.007 | $7.25 \times 10^{-3}$ |
| 14 | chr1:103573700-103573772 | 2 | cg26436330; cg20847625 | *COL11A1* | 0.01 | $6.33 \times 10^{-4}$ |
| 24 | chr:1045495981-45496216 | 3 | cg18382353; cg16512882; cg15078013 | *C10orf25*; *ZNF22* | -0.02 | $8.49 \times 10^{-4}$ |
| 9 to 14 | chr3:52569053-52569169 | 3 | cg18337363; cg08365687; cg13284614 | *NT5DC2*; *LOC440957* | -0.009 | $7.35 \times 10^{-3}$ |
| 14 to 24** | chr17:2699706-2699718 | 2 | cg05890550; cg25373595 | *RAP1GAP2* | 0.02 | $6.53 \times 10^{-5}$ |

*Note. Bonf.-adjust = Bonferroni-adjusted. A negative effect size (-) indicates hypermethylation associated with faster latency (for cross-sectional measurements) or latency becoming slower overtime (for change scores). \*\*This row details the topmost significant DMR to associate with 14- to 24-month latency change (see SM 4 Table 2 for a summary of all DMRs significantly associated with PLR latency).*

### PLR amplitude

No probes were found to be significantly associated with PLR amplitude after stringent epigenome-wide multiple comparisons corrections (*p* < 2.4 × 10^−7^). Several differentially methylated PLR amplitude-associated probes reached the discovery p-value threshold *p* < 5 × 10^−5^ (See full list in SM 4 Table 1). The region analyses revealed 17 DMRs significantly associated with PLR amplitude with the topmost significantly associate DMR per phenotype listed in Table 4 (full list in SM 4 Table 2).

**Table 4.** Summary of the topmost significantly associated DMRs for each PLR amplitude phenotype.

| Amplitude phenotype (months) | Genomic locations (hg19) | N probes | Probes | Illumina gene annotation | Effect size ($\beta$) | Bonf.-adjust p-value |
| --- | --- | --- | --- | --- | --- | --- |
| 9 | chr13:103452556-103453215 | 4 | cg06518779; cg21251000; cg15186648; cg15193473 | *BIVM*;<br>*KDELC1* | 0.04 | $2.13 \times 10^{-4}$ |
| 14 | chr11:111249659-111250201 | 5 | cg19126910; cg17390301; cg24049888; cg18316498; cg11362935 | *POU2AF1* | -0.03 | $2.22 \times 10^{-2}$ |
| 24 | chr5:77253833-77253990 | 3 | cg25051331; cg09048251; cg07595776 | | -0.08 | $8.23 \times 10^{-4}$ |
| 9 to 14 | chr10:102295134-102295549 | 5 | cg07690778; cg26303175; cg07080220; cg08314679; cg07510080 | *HIF1AN* | 0.07 | $2.67 \times 10^{-3}$ |
| 14 to 24 | chr15:89786761-89787223 | 4 | cg22813622; cg01741397; cg15769724; cg06870609 | *FANCI* | -0.04 | $1.22 \times 10^{-3}$ |

*Note. Bonf.-adjust = Bonferroni-adjusted. A negative effect size (-) indicates hypermethylation associated with smaller constriction amplitude (for cross-sectional measurements) or constriction becoming larger overtime (for change scores).*

### Downstream Exploratory analysis

Gene ontology (GO) analysis was conducted for each phenotype using genes annotated with probes within the top 1000 EWAS p-values, and probes located within the significantly associated DMRs. Multiple significantly enriched GO terms were identified (Table 5). SM 5 Table 2 lists all significantly enriched GO terms.

**Table 5.** Summary of the number of significantly enriched GO terms and the most significant term for each PLR phenotype.

| Phenotype (month) | N. sig. enriched GO terms | Most significantly enriched GO term<br>Description (GO number) | Fold Enrichment | FDR p-value |
| --- | --- | --- | --- | --- |
| **<u>PLR latency</u>** |  |  |  |  |
| 9 | 12 | Homophilic cell adhesion via plasma membrane adhesion molecules (GO:0007156) | 4.2 | $3.17 \times 10^{-7}$ |
| 14 | 34 | Negative regulation of developmental process (GO:0051093) | 1.84 | $6.95 \times 10^{-3}$ |
| 24 | 0 | - | - | - |
| 9 to 14 | 16 | Homophilic cell adhesion via plasma membrane adhesion molecules (GO:0007156) | 6.62 | $8.27 \times 10^{-22}$ |
| 14 to 24 | 16 | Regulation of cellular component organization (GO:0051128) | 1.5 | $5.00 \times 10^{-3}$ |
| **<u>PLR amplitude</u>** |  |  |  |  |
| 9 | 0 | - | - | - |
| 14 | 73 | Regulation of glucuronosyltransferase activity (GO:1904223) | 24.41 | $3.86 \times 10^{-8}$ |
| 24 | 0 | - | - | - |
| 9 to 14 | 22 | System development (GO:0048731) | 1.41 | $1.30 \times 10^{-3}$ |
| 14 to 24 | 0 | - | - | - |

*Note. sig. = significant after False discovery rate (FDR) adjustment.*

For both PLR components, no probes above discovery p-value threshold, nor probes located in any of the significantly associated DMRs, were listed in the MRC-IEU EWAS catalogue (50). None of the stringently significant probes were found to be annotated to genes listed in the SFARI database (51). Nineteen probes significantly associated with PLR at the discovery p-value threshold were annotated to 18 distinct genes listed in the SFARI gene. Notably two SFARI genes with ‘high confidence’ associations were annotated with probes identified to significantly associate with latency at 14 months (cg01123282; *NR4A2*) and latency at 24 months (cg02230180; *HNRNPU*). A full list of probes annotated to SFARI genes are summarised in SM 5 Table 3.

Three of the identified significantly associated DMRs contained probes annotated to three distinct SFARI genes, as summarised in Table 6.

**Table 6.** Summary of significantly associated DMRs annotated to genes previously linked to autism.

| DMR genomic locations (hg19) | Probes | Associated phenotype | Gene | SFARI gene-score* |
| --- | --- | --- | --- | --- |
| chr11:19372012-19372234 | cg23330281;<br>cg24137774;<br>cg25909885 | Latency 14 to 24 month | *NAV2* | 2<br>(strong candidate) |
| chr7:2968559-2968595 | cg22989995;<br>cg23770265 | Amplitude 24 month | *CARD11* | 2<br>(strong candidate) |
| chr13:38444227-38444490 | cg16409955;<br>cg23021771 | Amplitude 9 to 14 month | *TRPC4* | 3<br>(suggestive evidence) |

*Note: \*SFARI gene-scores is a ranking system that reflect the overall strength of evidence of the gene’s relevance to autism (54).*

## Discussion

The infant pupillary light reflex (PLR) is a candidate early intermediate phenotype for autism, offering promise as a biomarker and tool to investigate mechanisms underlying autism emergence. We analysed epigenome-wide probe– and region-level differential DNA methylation (DNAm) with variability in PLR onset latency and constriction amplitude between 9-months and 2 years. We then performed functional exploratory analyses of PLR-associated probes and regions.

We identified four novel epigenome-wide-significant differentially methylated probes associated with three PLR latency phenotypes. Specifically, hyper-methylation of cg05148717 and cg22367466 were associated with faster 14-month latency; hypermethylation of cg09732535 was associated with slower 24-month latency; and hyper-methylation of cg15130433 was associated with PLR latency becoming faster between 14– and 24-months. Thirteen differentially methylated regions were associated with latency, particularly the 14-to 24-months change. These findings provide novel evidence highlighting the role of differential epigenetic signature in the variability of early PLR onset latency.

PLR latency and DNAm have been linked to the emergence of autism characteristics (5,13,28). This, alongside the current findings, implicates PLR latency as a potential mechanistic bridge linking DNAm to autism. To explore this, we examined whether associated probes or regions were listed in the SFARI database (51) or the EWAS catalogue (50). While none of the stringently significant probes or regions had prior links to autism, as reported in these two databases, several discovery significant probes across the five latency EWASs were annotated to autism susceptibility genes. Among the most compelling genes were *NR4A2* and *HNRNPU*, listed as ‘high confidence’ autism-linked genes in the SFARI database given the multiple separate reports identifying mutations in each of these genes in individuals with autism (52–56). Probe cg01123282, annotated to intron 1 of *NR4A2*, showed hypermethylation associated with slower latency at 14 months. The hypermethylation of probe cg02230180, annotated to intron 1 of *HNRNPU*, was associated with faster PLR latency at 24 months. Both genes are involved in neuronal functioning and development; *NR4A2* is implicated in dopaminergic neural cell differentiation (57), and *HNRNPU* is linked to synaptic function (58) and neurogenesis (59). Echoing the involvement of epigenetic regulated neurodevelopmental genes in the modulation of PLR latency, gene ontology (GO) analysis indicated probes associated with each of the PLR latency phenotypes, except for cross-sectional 24-month, were annotated to genes involved in neurodevelopmental or cellular organisation biological processes. These included neurogenesis (GO:0022008), nervous system development (GO:0007399), negative regulation of developmental process (GO:0051093), and regulation of cellular component organization (GO:0051128). These findings are notable given previous evidence pointing to the contributing role of DNAm in the perturbations of neurodevelopmental processes in autism (18). Collectively, the current findings indicate the epigenetic regulation of PLR latency, supports PLR as a candidate biomarker, and underscores the potential perturbations to key neurodevelopmental processes as candidate biological mechanisms linking DNAm, PLR latency development and the early emergence of autism characteristics.

In contrast to PLR latency, no individual probes were significantly associated with PLR amplitude after stringent epigenome-wide multiple testing corrections. This difference between the two PLR components likely reflects distinct biological regulatory mechanisms. Although the pupil reflex is primarily regulated by the core four-neuron reflex circuity (7), a growing body of evidence indicates PLR amplitude is influenced by additional top-down modulation from higher-order cognitive processes, such as emotional regulation and executive control (60–63). GO analysis in the current paper further indicates PLR amplitude regulation to be complex, having identified amplitude-associated probes annotated to genes covering a more diffuse and complex biological network, relative to PLR latency. Amplitude-related GO terms included developmental processes (system development, GO:0048731) and cellular organization (homophilic cell adhesion via plasma membrane adhesion molecules, GO:0007156), like PLR latency, but also included a larger proportion of terms like cellular signalling (regulation of signalling, GO:0023051), metabolic regulation (regulation of glucuronosyltransferase activity, GO:1904223), and stress response (regulation of response to stimulus, GO:0048583). The top-down regulatory landscape of amplitude, alongside this relatively broader biological enrichment, likely weakens the impact of single-probe differential DNAm. In contrast, if latency is regulated by a narrower and predominantly bottom-up neurobiological network, then probe-level differential DNAm in this network is likely more penetrant and detectable after stringent multiple comparison corrections.

Indeed, at a discovery threshold, DNAm of multiple probes was significantly associated with PLR amplitude, as were several regions. PLR amplitude has previously been linked to autism (5,13,14), and although no discovery significant probes nor probes within significantly associated regions were found to have direct links to autism according to the EWAS catalogue (50), twelve genes annotated with amplitude-associated probes were in the SFARI database (51). Nine of these were categorised as ‘strong candidates’ for autism. These autism-linked genes reiterate the diversity of the biological pathways connected to PLR amplitude highlighted in the GO analysis, with gene functions ranging from immunity (*CARD11)* (57), to metabolic pathways and circadian regulation (*CSNK2B*) (57,64), to glycoprotein production (*GALNT10)* (57), and calcium signalling pathways (*CACNA1D*) (57). Several of the identified autism-linked genes also play a role in neurodevelopment (57).

These findings provide initial support for PLR as a potential biomarker for autism and suggest the involvement of differential DNA methylation across a relatively broad network in linking early PLR amplitude to the emergence of autism characteristics.

### Limitations and future investigations

Our findings must be interpreted in the light of certain limitations. Our sample of male infants was enriched for increased familial likelihood for autism with over 80% of the sample being the younger siblings of an individual with clinical autism. This limits the generalisability to other populations, such as females or individuals with different autism liabilities (e.g., syndromic autism). However, studying DNAm in a homogeneous sample, with DNA and phenotypic measures collected within a tight timeframe, may enhance power, as such homogeneity likely reduces signal-to-noise ratios, leading to larger effect sizes (28,65). Supporting this, although the sample size is relatively small, four epigenome-wide significantly associated probes were identified. Larger samples may reveal additional associations between PLR latency and DNAm, as well as offering deeper insights into the relationship between PLR amplitude, epigenetics and autism.

A second potential limitation is that quantifying DNAm of PLR in the reflex neurocircuitry tissue, instead of buccal cells, may yield more specific results related to PLR. However, buccal cells, being more accessible, allow for a larger sample size from live individuals that are deeply phenotyped over time (like our current sample), and more suitable as a biomarker than post-mortem brain tissue. Further, empirical evidence shows that DNAm patterns in various brain regions are more similar to saliva samples (which contain many buccal cells) than blood samples (66), and we accounted for variance attributable to differences in cell type proportions by including the proportion of common cell types in buccal samples as a covariate in our analysis.

Another limitation is that while the Illumina 450k array is a reliable tool for quantifying numerous variable DNAm sites, it only covers 2% of all DNAm sites across the genome (67). Limited coverage may hinder conclusions as probes related to PLR or autism may be unmeasured. Additionally, although we aimed to assess the association of PLR with early altered DNAm (at 9 months), this window for observing the role of DNAm in regulating PLR is narrow. Investigating DNAm at other timepoints or longitudinal changes in DNAm across infancy may offer complementary insights.

### Conclusions

In conclusion, this is the first study, to our knowledge, that has investigated the involvement of DNAm in the variability of infant PLR, a candidate early intermediate phenotype and candidate biomarker for autism. We identified DNAm of four probes to significantly associate, after epigenome-wide multiple comparison corrections (*p < 2.4 × 10^−7^*), with three PLR latency phenotypes over the 14-to 24-month period, and 13 differentially methylated regions were associated with latency, particularly the 14-to 24-months change. No probes were found to associate with PLR constriction amplitude after stringent multiple corrections. At discovery significance threshold, several probes were found to associate with both PLR components, as did multiple regions – some of which were annotated to autism susceptibility genes. Gene ontology and functional exploration pointed to possible biological mechanisms localised to nervous system and developmental processes for both PLR parameters but suggests amplitude to have a more complex biological underpinning. Future investigations should further unpick the functional implications of the identified differential methylation, as well as tackle limitations of the current study by replication with larger sample sizes and within other sub-groups of autism (e.g., females, or individuals from simplex families), exploring the association of longitudinal DNAm changes, DNAm in diverse tissues (e.g., neural regions) and with more comprehensive assays of DNAm (e.g., Illumina EPIC microarray or future iterations, or whole epigenome long-read sequencing).

## Funding

This work was funded by the Simons Foundation Autism Research Initiative (SFARI) Pilot Award grant no. 511504; European Union’s Horizon 2020 research and innovation programme under the Marie Skłodowska-Curie grant no.642996 (BRAINVIEW). Data collection was funded by MRC Programme [grant numbers G0701484 and MR/K021389/1], the BASIS funding consortium led by Autistica (www.basisnetwork.org), EU-AIMS (the Innovative Medicines Initiative joint undertaking grant agreement number 115300, resources of which are composed of financial contributions from the European Union’s Seventh Framework Programme [grant number FP7/2007-2013] and EFPIA companies’ in-kind contribution) and AIMS-2-TRIALS (Innovative Medicines Initiative 2 Joint Undertaking under grant agreement number 777394. This Joint Undertaking receives support from the European Union’s Horizon 2020 research and innovation programme and EFPIA and AUTISM SPEAKS, Autistica, SFARI). Epigenetic data generation was funded by MRC Centenary Award to Chloe C. Y. Wong.

Chloe C. Y. Wong was supported by the National Institute for Health and Care Research (NIHR) Maudsley Biomedical Research Centre at South London and Maudsley NHS Foundation Trust and King’s College London [NIHR203318]. The views expressed are those of the authors and not necessarily those of the NIHR, the Department of Health, or King’s College London. Laurel A. Fish was supported by the SGDP PhD Studentship award.

## Ethics

NHS Research Ethics Committees granted ethical approval (08/H0718/76 [BASIS] and 06/MRE02/73 [DNA collection, extraction and analysis]). Informed written consent was provided by the parent(s).

## Acknowledgement

We heartfully thank all the families who took part in our research. We also thank The BASIS Team members listed here in alphabetical order contributed to the design and/or data collection of the longitudinal BASIS study: Baron-Cohen S., Blasi A., Cheung C., Davies K., Elsabbagh M., Fernandes J., Gammer I., Ganea N., Gliga T., Guiraud J., Hendry A., Liersch U., Liew M., Maris H., O’Hara L., Pickles A., Ribeiro H., Salomone E., Taylor, C., Tucker L., Tye C., Wass S.

The authors acknowledge the contribution and use of the CREATE high-937 performance computing cluster at King’s College London (King’s College London (2022). King’s Computational Research, Engineering and Technology Environment (CREATE), https://doi.org/10.18742/rnvf-m076.).

## Notes

### Competing Interest Statement

The authors have declared no competing interest.

