## Supplemental Material for "Epigenome-wide analysis reveals novel DNA methylation signatures significantly associated with the infant pupillary light reflex, a candidate intermediate phenotype for autism"

**Supplementary Material**

[SM 4.2: EWAS probes at discovery p-value threshold (p < 5 × 10^-5^) or above.	14](#_Toc187226000)

### SM 1: Study cohort and procedure

The British Autism Study of Infant Siblings (BASIS; [www.basisnetwork.org](http://www.basisnetwork.org/)) is a prospective longitudinal study aiming to track the emergence of autism characteristics in a sample of children with about 20% increased likelihood for receiving a later autism diagnosis due to having a first degree relative with a diagnosis of autism(1). Measurements are taken at 9 months, 14 months, 24 months and 3 years using a battery of parent-report questionnaires, eye-tracking and EEG experimental tasks, standardised behavioural assessments and clinical assessments. Infants were born full-term (36+ weeks) and had no known genetic syndromes or visual, auditory, or other disabilities.

The sample for the current paper participated in the second phase of the BASIS and had buccal samples taken for epigenetic investigations at 9 months. SM 1 Table 1 summarises the sample characteristics.

SM 1 Table 1. Summary of sample for each EWAS including total number of participants (and number of those with increased and low familial likelihood for autism), average age and average PLR measure.

| Phenotype | N  (IL, LL)^1^ | Age in months  Mean (SD) | PLR  Mean (SD)^2^ |
| --- | --- | --- | --- |
| **Latency (ms)** |  |  |  |
| 9 months | 48  (40, 8) | 9.07  (0.85) | 329.18 (40.05) |
| 14 months | 47  (38, 9) | 15.44  (1.02) | 317.54 (41.05) |
| 24 months | 40  (32, 8) | 26.22  (2.04) | 312.41 (29.38) |
| 9 to 14 months | 44  (36, 8) | - | -15.45 (35.56) |
| 14 to 24 months | 37  (29, 8) | - | 0.93  (39.78) |
| **Amplitude (%)** |  |  |  |
| 9 months | 46  (38, 8) | 9.06  (0.85) | 32.89  (7.71) |
| 14 months | 47  (38, 9) | 15.44  (1.02) | 36.16  (8.27) |
| 24 months | 37  (29, 8) | 26.15  (2.09) | 28.12  (7.81) |
| 9 to 14 months | 42  (34, 8) | - | 2.90  (7.21) |
| 14 to 24 months | 34  (26, 8) | - | -8.32  (8.81) |

*Note: ^1^Sample included in EWAS split by autism diagnosis likelihood (LL = low familial likelihood; IL = increased familial likelihood). ^2^PLR values are the raw (non-residual) values.*

### SM 2: PLR processing and covariates

#### SM 2.1: PLR Processing

The PLR processing pipeline implement is the same as that described previously (2,3) with aspects of the pipeline optimised using observations of the current dataset, for parameter identification (e.g. latency and maximum constriction window) and attrition (e.g. later exclusion criterion). Pupils were measured to be at an average distance of 59.54 cm (59.15 cm–59.95 cm) during the PLR. Following previous work(2), to control for differing PLR inducing light levels across the white slide in stimulus one, only trials where the first look was within the centre 5% of the white slide (see SM 2 Figure 2) were included. First, an automated cleaning algorithm was applied to data using R (4) and R-studio (5). This algorithm involved: the removal of measurement outside the range of 1-10mm diameters; linear interpolation of short “flicker” gaps (<7 samples) with <0.2mm diameter change; removal of samples with large diameter changes from the previous sample (>0.3mm); removal of short segments of data (<6 samples); additional linear interpolation; resampling data to 300hz (6) and application of a 25-point moving average filter (2) supported by visual inspection of data. This cleaning algorithm resulted in a continuous smoothed pupil trace that, compared to the raw pupil trace, had a higher resolution and fewer potential artefacts. Using this pupil trace, we calculated first (velocity) and second (acceleration) order derivatives, while applying 25-point moving average filters between each derivation to reduce noise amplification (6). Time windows for minimum and maximum constriction were determined by a series of optimisations investigations. The PLR latency was defined by the acceleration minima in the time interval 110-570ms to the stimuli onset. The baseline pupil size for each trial was defined as the average pupil size in a 100ms interval just before the latency time point. The amplitude of the PLR was calculated using the formula presented in Figure 1 in the main text, using the maximum constriction within the interval 170– 1450ms relative to latency onset.

All pupil traces were visually inspected for manual validation and were rejected based on two criteria: a) just the latency was incorrectly identified or b) both the latency and the maximum constriction was incorrectly identified. For example, if a pupil trace had a valid latency but invalid maximum constriction (e.g. the infant blinked after the PLR latency, but before maximum constriction), then the trace was accepted only for its valid latency and included in latency analysis only (not amplitude analysis). We conducted inter-rater reliability checks on the manual validation. A second independent validator scored 20% of trials. Unweighted Cohen’s Kappa Coefficient was used to assess the agreement between two manual validators’ judgement on whether PLR processing across the whole all measurement from the BASIS sample (phase 2 and phase 3) correctly extracted PLR latency or the maximum constriction (latency: κ = .86 [95% CI, .84 to .88], p < .001, constriction: κ = .84 [95% CI, .82 to .86], p < .001).

Once manual validation had been completed, those trials where either latency or both latency and maximum constriction points had been successfully identified were passed to the second pipeline of automated processing. For each trial, the eye (left or right) with the pupil trace that best correlated (parallel correlations according to p-value and correlation coefficient) with all other valid pupil traces for that participant at that assessment timepoint were extracted. If neither the left nor right eye’s trace significantly correlated with the remaining manually validated traces, then that trial was excluded.

Trials were excluded if: a) the latency window had more than 75% interpolated or missing data in original trace; b) the maximum constriction window had more than 75% interpolated or missing data in original trace; c) the amplitude was outside a range of 5% to 80% (thus biologically implausible); d) 100% of the window +/-40ms relative to latency or maximum constriction was interpolated data (+/-40ms being the size of the 25pt moving average window around the latency/maximum constriction point). Participants with fewer than 3 trials remaining after processing were excluded. The median latency and amplitude per individual per timepoint were calculated and included in models as dependent variables. The averaging of PLR parameters was conducted in line with findings (6) whereby authors concluded that for 60-Hz sampling rate, averaging latencies of multiple pupil light reflexes helped to improve the resolution of latency because the average latency was not constrained to a time grid of 16.7ms.

#### SM 2.2 PLR covariates

We considered the trials missingness (SM 2.2.1) and baseline pupil size (SM 2.2.2) as a potential covariate for our PLR measures.

##### SM 2.2.1 Missingness

The PLR, especially latency, is suspectable to number of trials included in the averaging(6). Trials could be missing either due to children not watching the stimuli (e.g., if they were distracted or inattentive during the protocol) or were excluded during PLR processing due to poor data quality. We calculated missingness (see SM 2 Table 1) as a percentage from of the total number of possible trials in the stimuli (32 for 9/14 months, 16 for 24 months). Missingness was averaged across trials within participant for each timepoint.

SM 2 Table 1. Mean percentage of trials missing during processing for each PLR parameter at each timepoint.

|  | Missing (%)  Mean (SD) | | |
| --- | --- | --- | --- |
|  | 9 months | 14 months | 24 months |
| Latency | 59.51 (13.54) | 58.98 (14.05) | 47.81 (20.73) |
| Amplitude | 62.09 (13.2) | 62.77 (14.33) | 55.41 (20.1) |

*Note: values are an average of individual’s missingness percentages. SD = standard deviation*

##### SM 2.2.2 Baseline pupil size

The size and strength of the PLR is potentially influenced by the size of pupil prior to the PLR(7). We calculated baseline pupil size (see SM 2 Table 2) for each trial as the average pupil diameter between 100ms before PLR latency and the PLR latency. Baseline pupil size was averaged across trials within participant for each timepoint.

SM 2 Table 2. Mean baseline pupil for each PLR parameter for each timepoint.

|  | Baseline pupil size (mm)  Mean (SD) | | |
| --- | --- | --- | --- |
|  | 9 months | 14 months | 24 months |
| Latency | 3.56 (0.41) | 3.62 (0.41) | 3.66 (0.39) |
| Amplitude | 3.55 (0.41) | 3.63 (0.41) | 3.69 (0.36) |

*Note: values are an average of individual’s baseline. SD = standard deviation*

##### SM 2.2.3. Accounting for covariates in PLR variables

To account for the variance in the PLR parameters (at each timepoint: 9, 14, and 24 months) attributable to the covariates, we conducted linear multiple regression models with PLR (latency or amplitude) as the dependent variable and covariates (missingness and baseline pupil size) as the predictor variables. Residuals from these analyses were extracted and used in all subsequent analysis as the PLR variables.

### SM 3: DNA Methylation data processing

#### SM 3.1. DNA Methylation level

Processing of DNA methylation resulted in 402,971 probes with methylation level data at each probe expressed as a 'beta' value (β) ranging from 0 (no methylation) to 1 (complete methylation) as illustrated in SM 3 Figure 1.

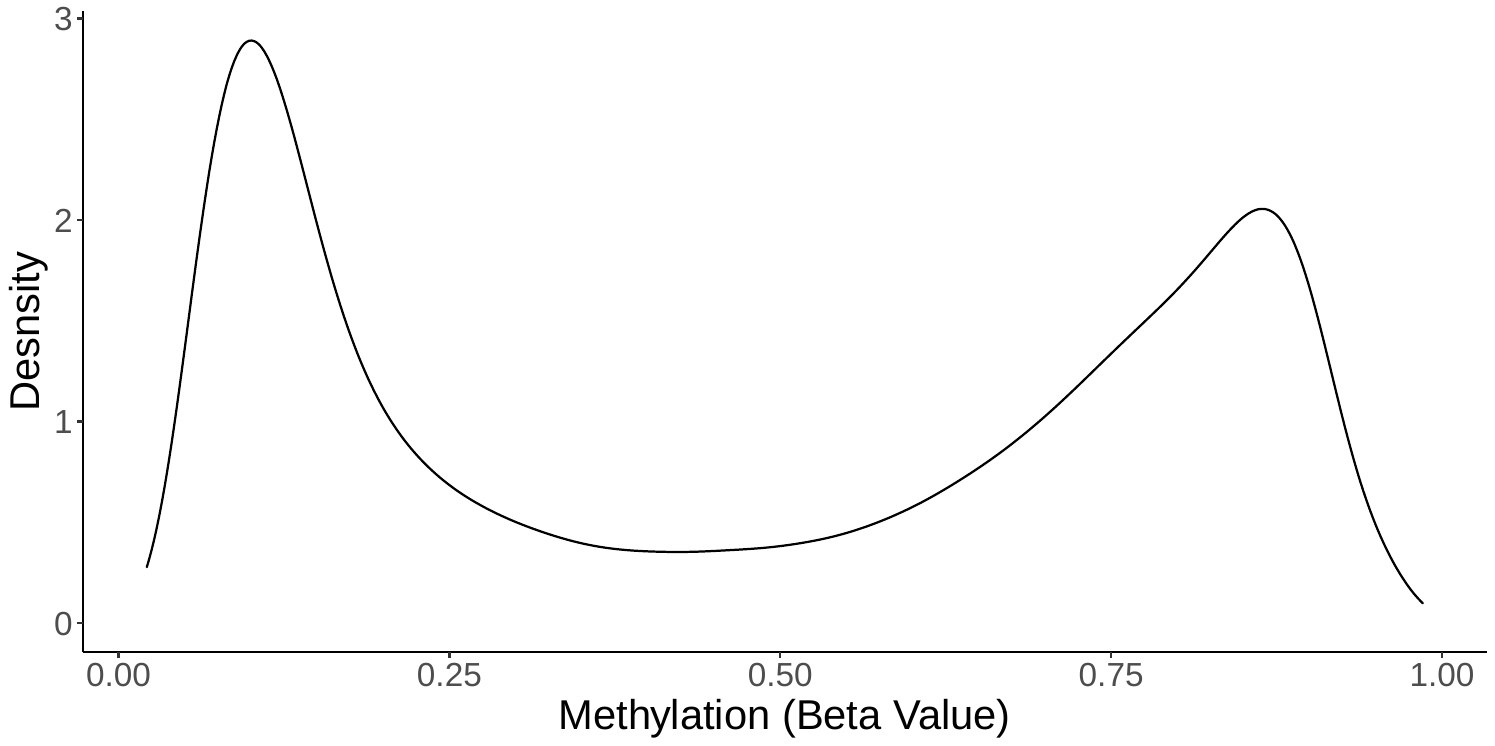

SM 3 Figure 1. Density plot of the distribution of β-values across sample.

#### SM 3.2: Location-based principal component for Epigenome-wide Analysis (EWAS)

To account for sources of unwanted variance attributable to population stratification in the processed DNAm beta values, we conducted a location-based principal components analysis and included the outputted 10 principal components as covariates in the EWAS model. This location-based principal component analysis(8) was computed using CpG sites within 50 base pairs of Single Nucleotide Polymorphisms (SNPs) with minor allele frequency > 0.01 reported in the 1000 Genomes Project (9). Restricting the PCA to CpG sites located near such SNPs narrows the focus of methylation variants that are better proxies of ancestry(8).

#### SM 3.3: Cell-type principal component for Epigenome-wide Analysis (EWAS)

To account for unwanted variance arising from cell-type signals in the processed DNA methylation beta values, we performed an analysis using EPiDISH(10). This analysis estimates methylation levels for nine common cell types found in buccal samples, allowing us to incorporate these estimates as covariates in our epigenome-wide association study (EWAS) model. Cell types included epithelial cells, fibroblasts, B cells, CD4+T cells, monocytes, eosinophils, neutrophils, CD8+ T cells, natural killer cells.

In our EWAS model, we included all estimated cell-type proportion as covariates, except for CD8+ T cells, which exhibited zero variance, and natural killer cells, which caused singularity in the EWAS model as it was highly correlated with B cell proportion (r = 0.98, p <0.001).

Including the estimated proportion for these cells improved model robustness and confounding effects related to cell-type composition of the buccal samples.

### SM 4: EWAS results

#### SM 4.1 Lambda inflation factor Q-Q plots and EWAS Manhattan plots for each EWAS

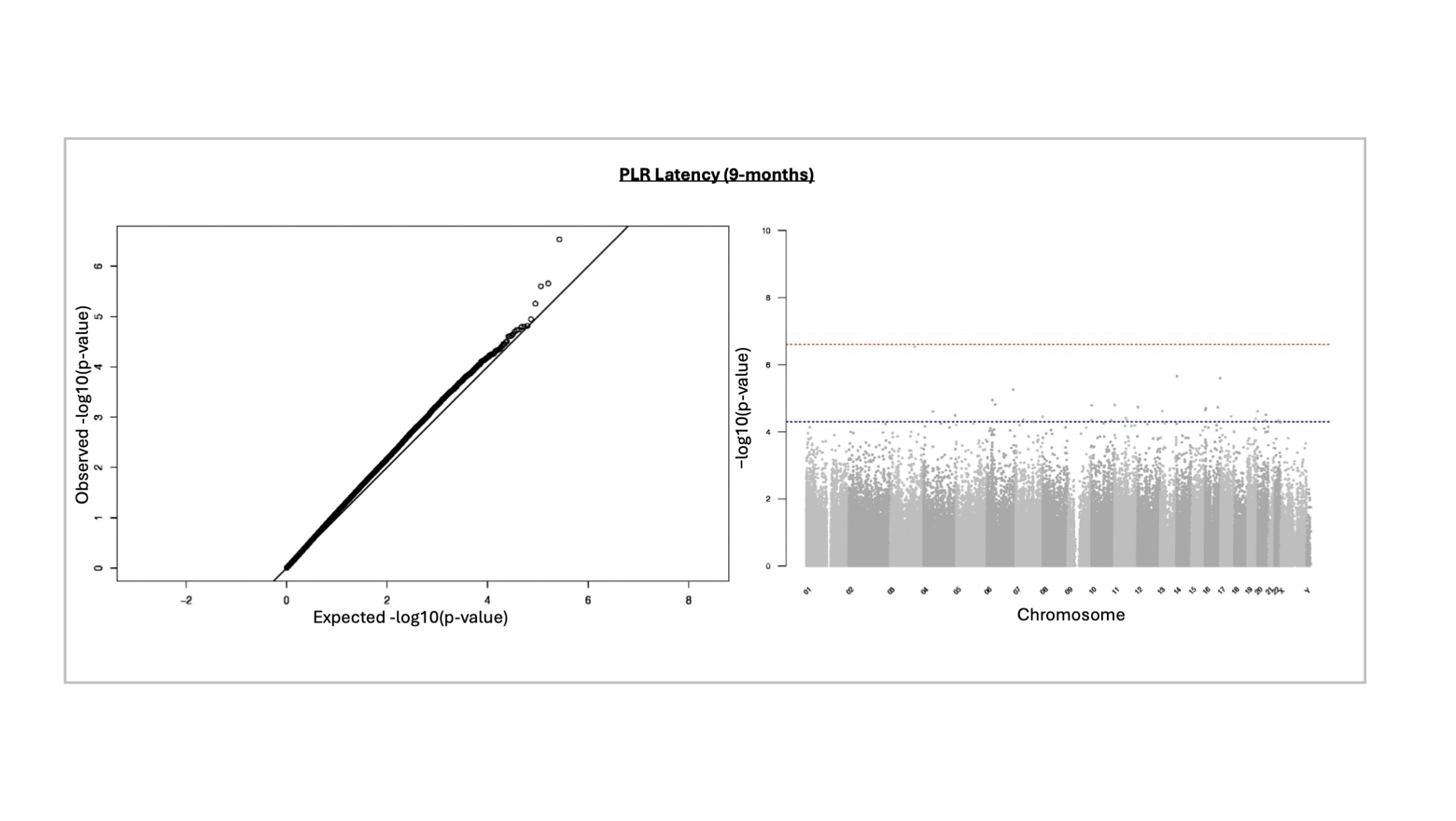

SM 4 Figure 1. Q-Q plot (left) and Manhattan plot (right) of results from 9-month PLR latency EWAS

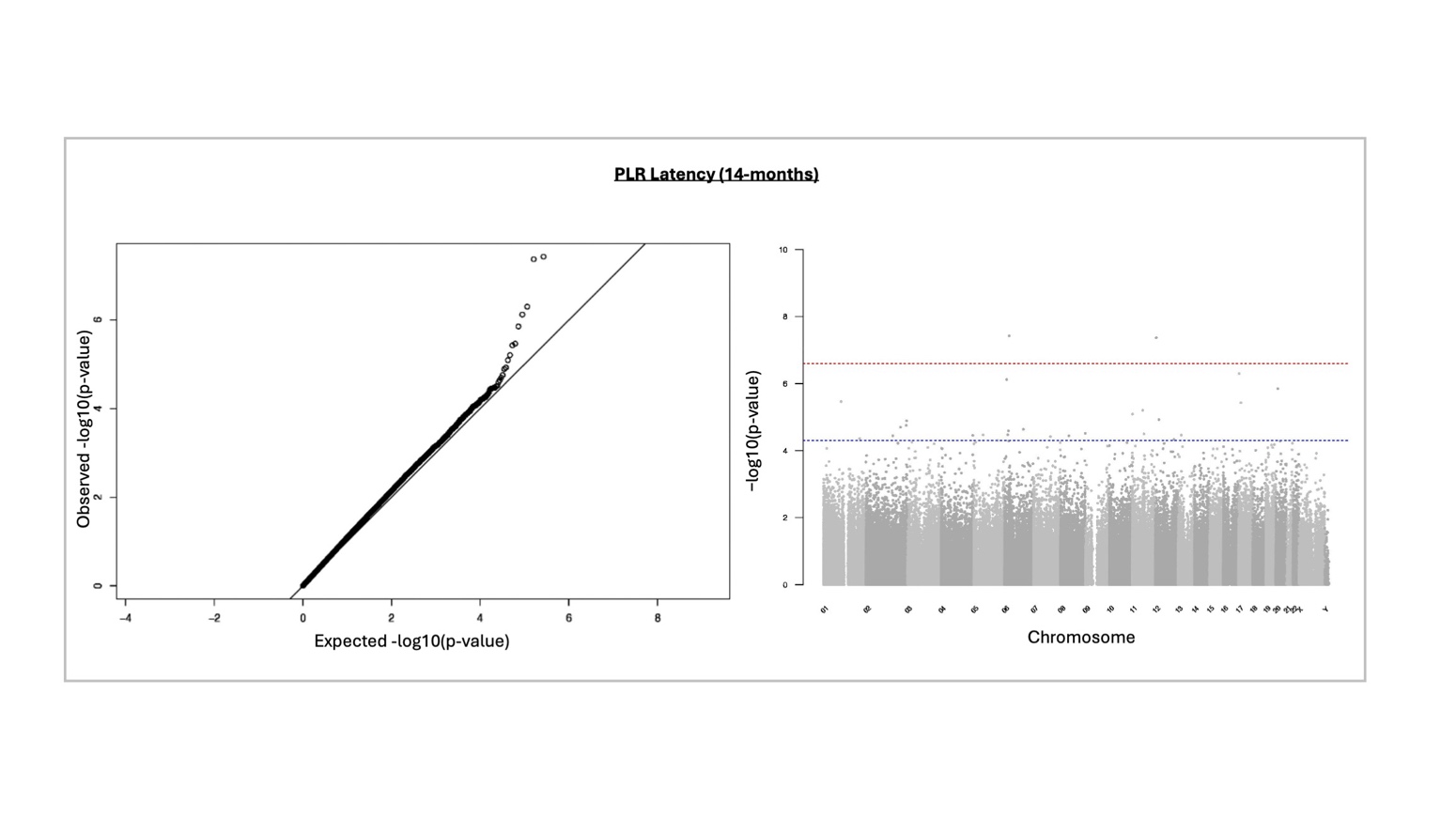

SM 4 Figure 2. Q-Q plot (left) and Manhattan plot (right) of results from 14-month PLR latency EWAS

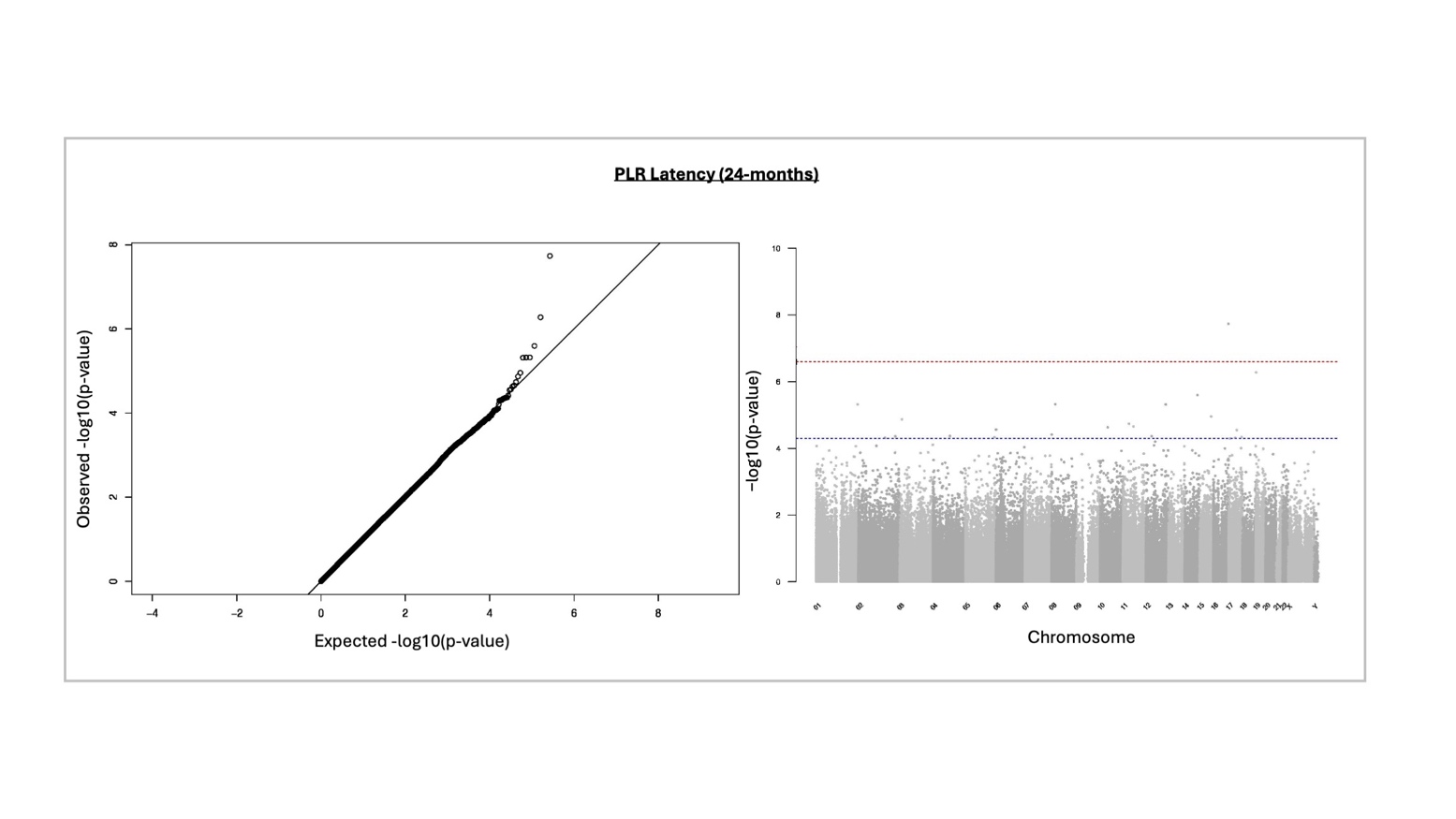

SM 4 Figure 3. Q-Q plot (left) and Manhattan plot (right) of results from 24-month PLR latency EWAS

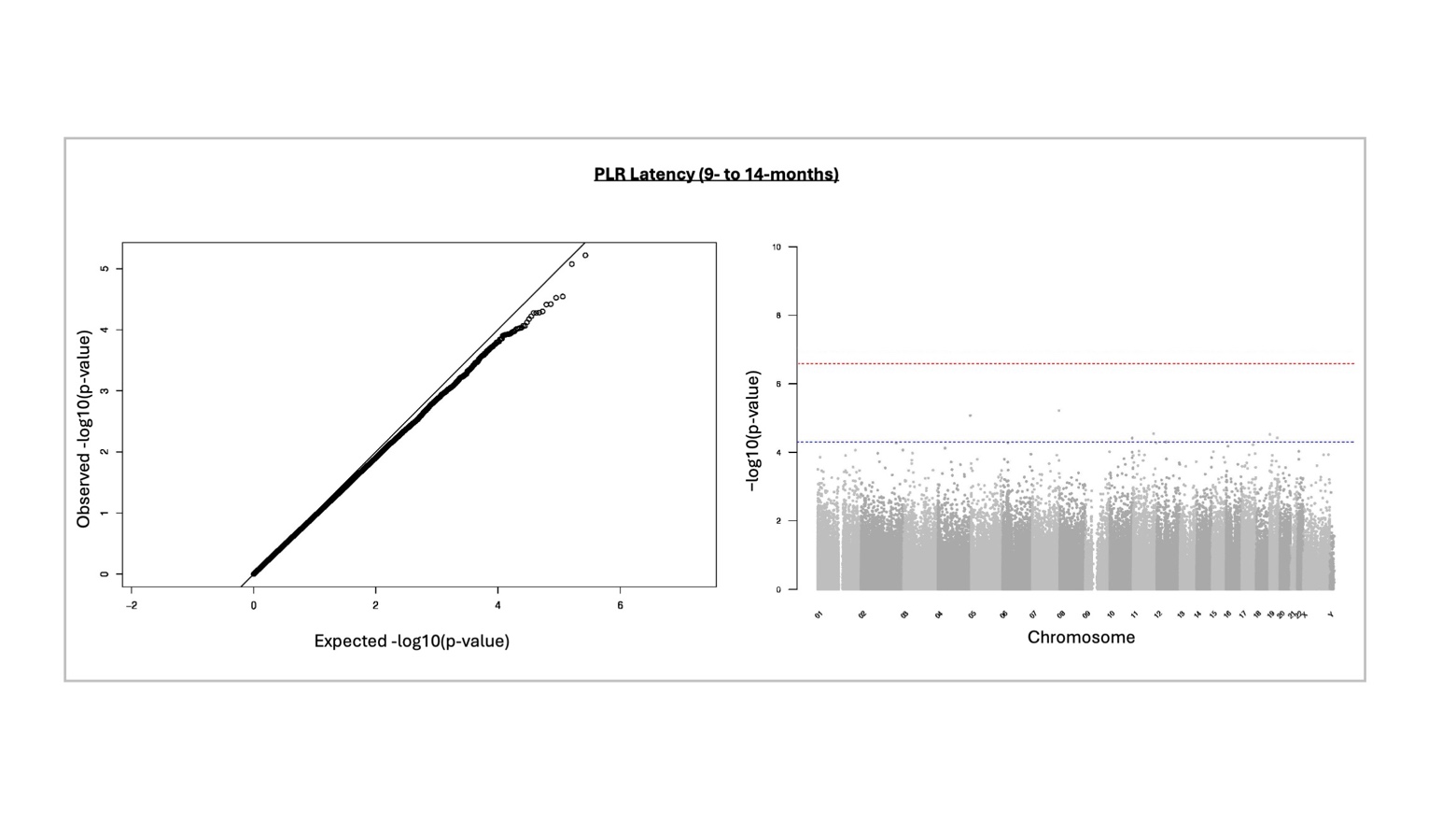

SM 4 Figure 4. Q-Q plot (left) and Manhattan plot (right) of results from 9- to 14-month PLR latency EWAS

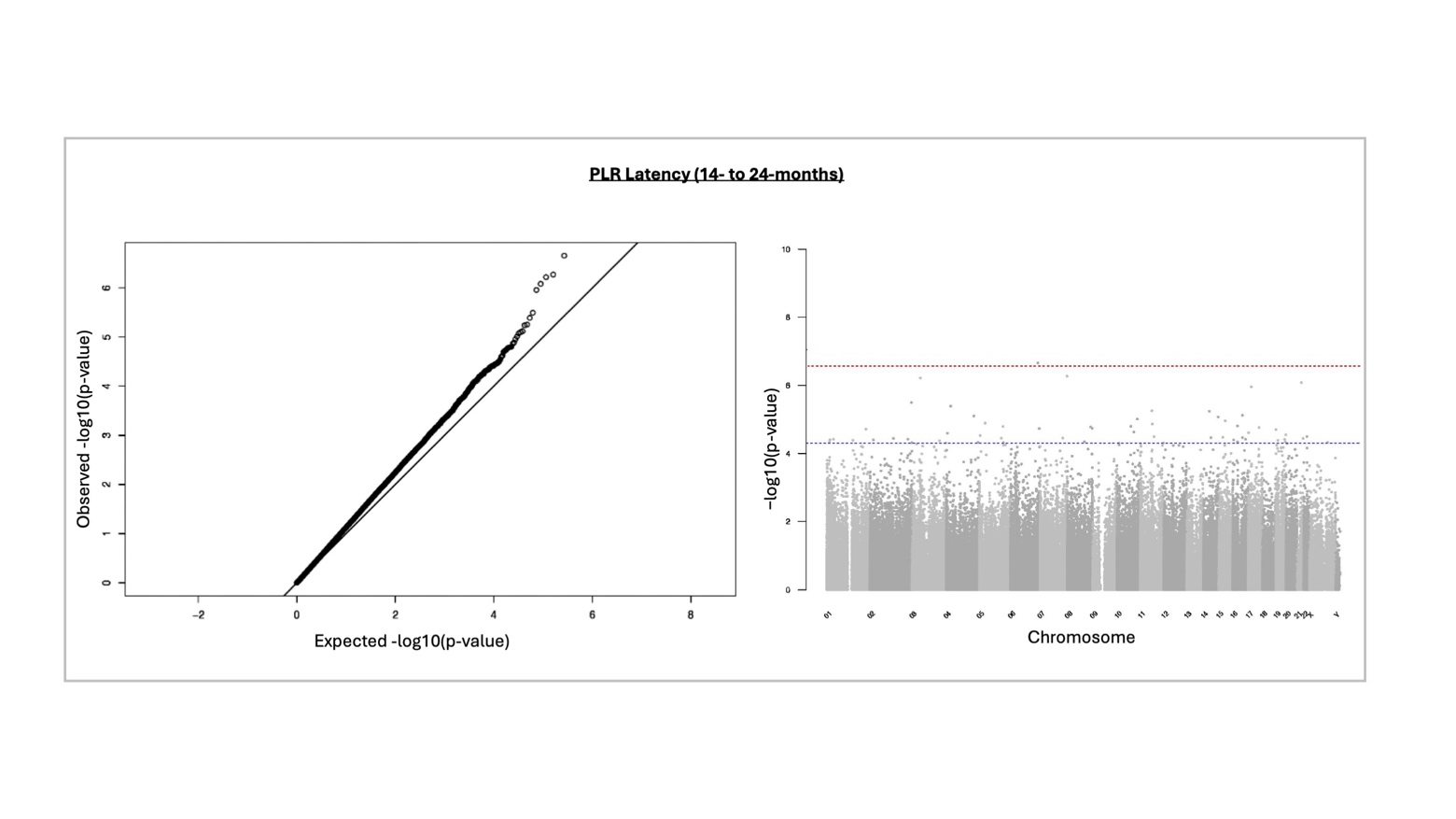

SM 4 Figure 5. Q-Q plot (left) and Manhattan plot (right) of results from 14- to 24-month PLR latency EWAS

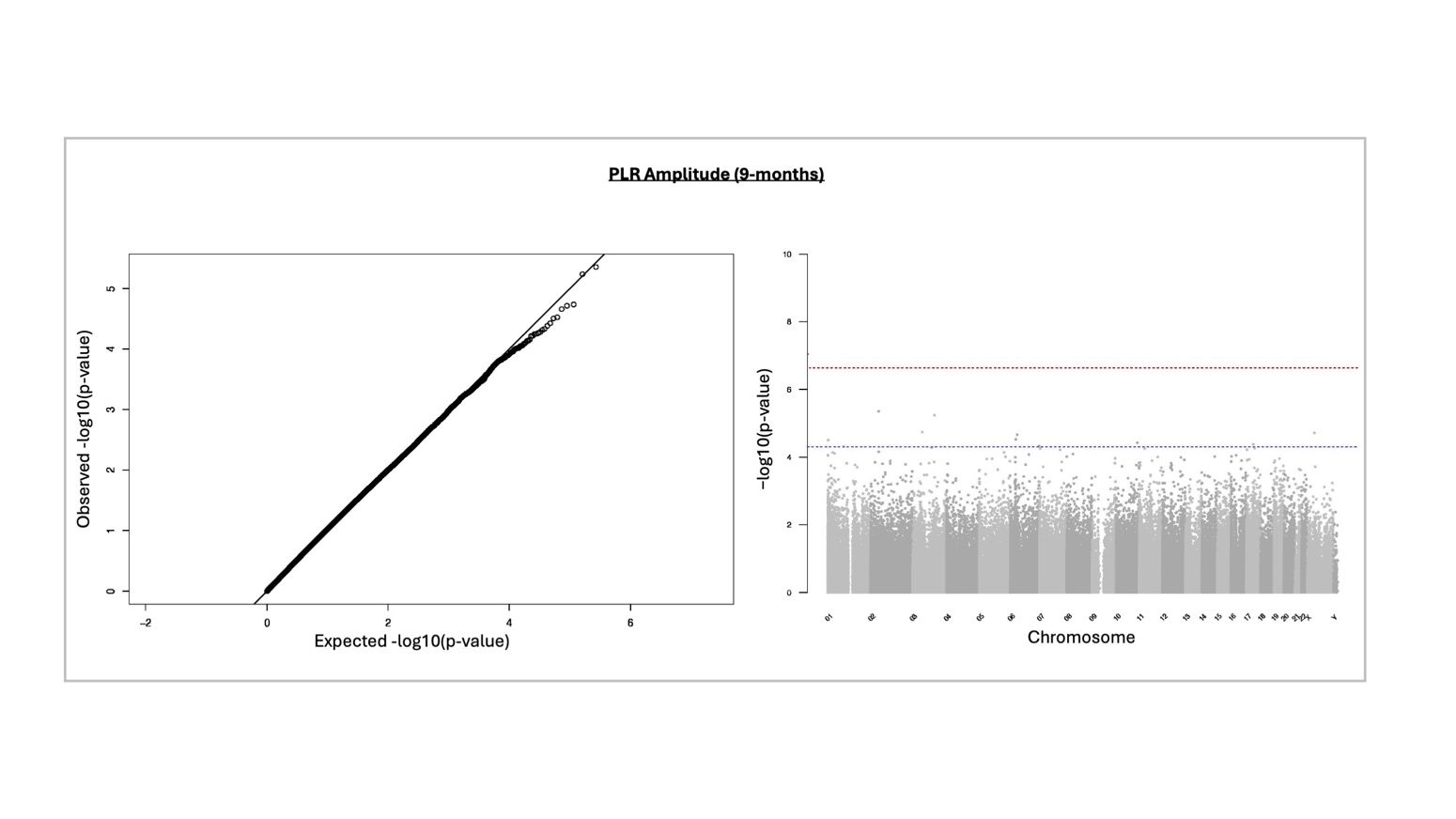

SM 4 Figure 6. Q-Q plot (left) and Manhattan plot (right) of results from 9-month PLR amplitude EWAS

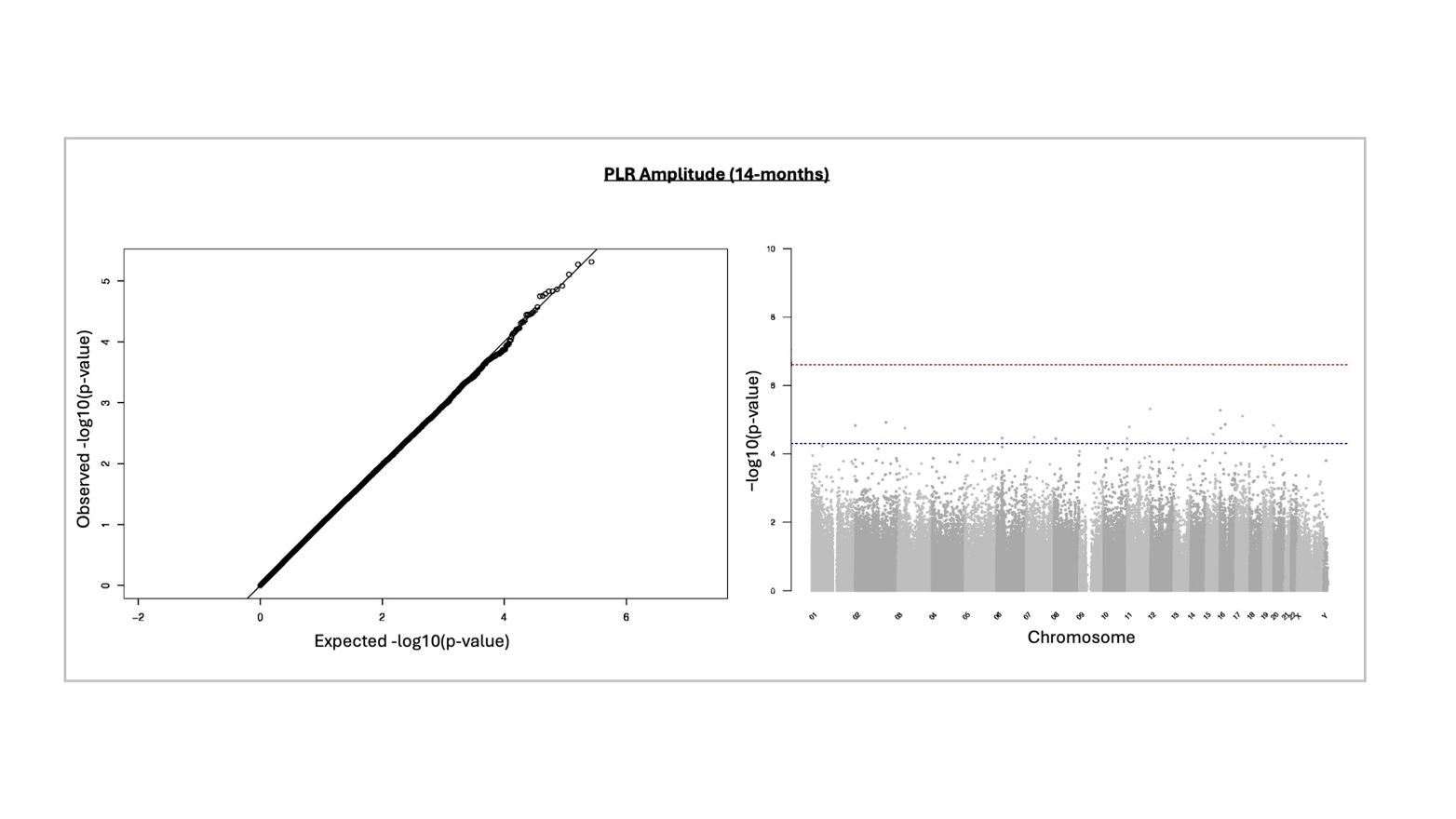

SM 4 Figure 7. Q-Q plot (left) and Manhattan plot (right) of results from 14-month PLR amplitude EWAS

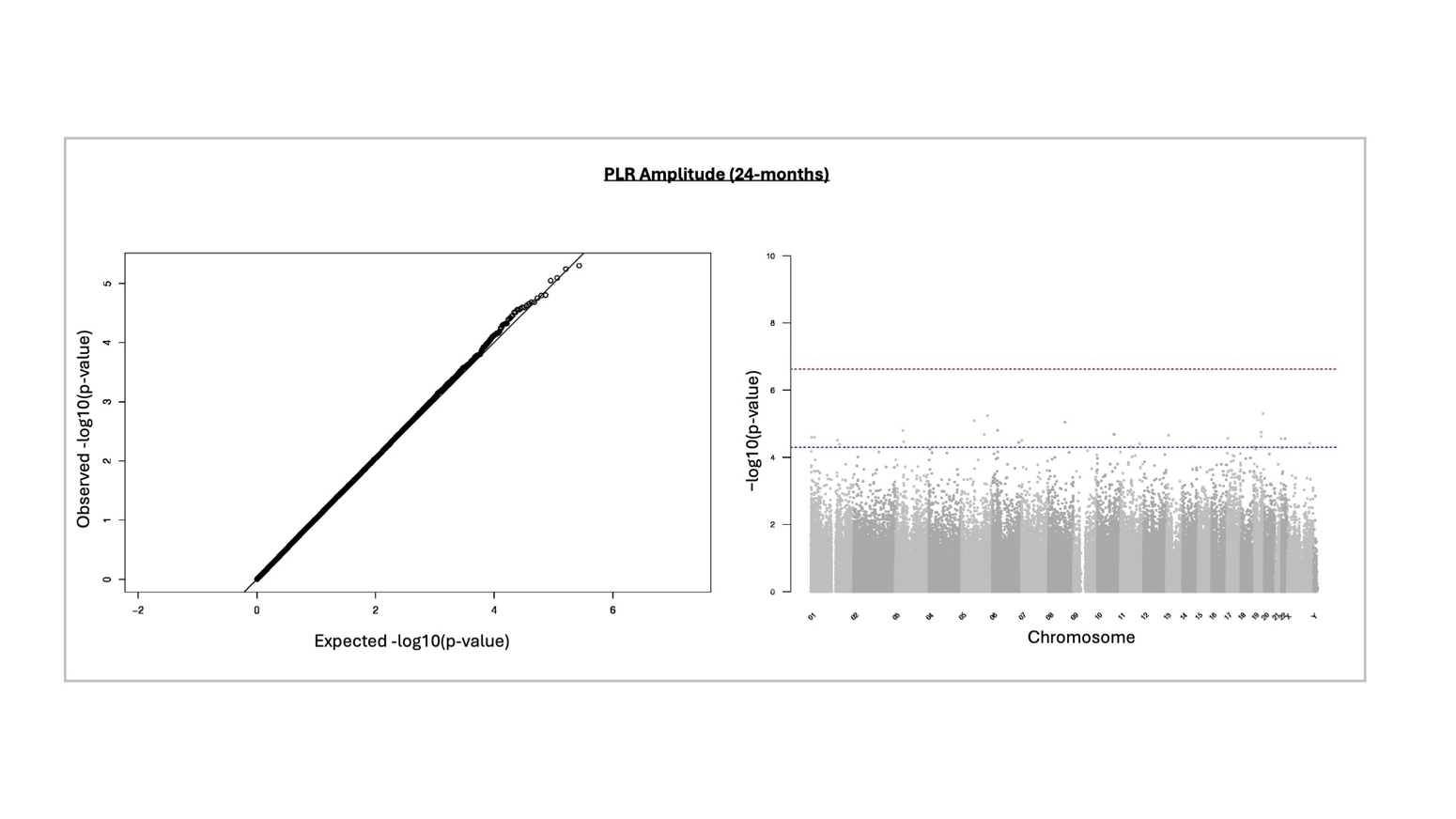

SM 4 Figure 8. Q-Q plot (left) and Manhattan plot (right) of results from 24-month PLR amplitude EWAS

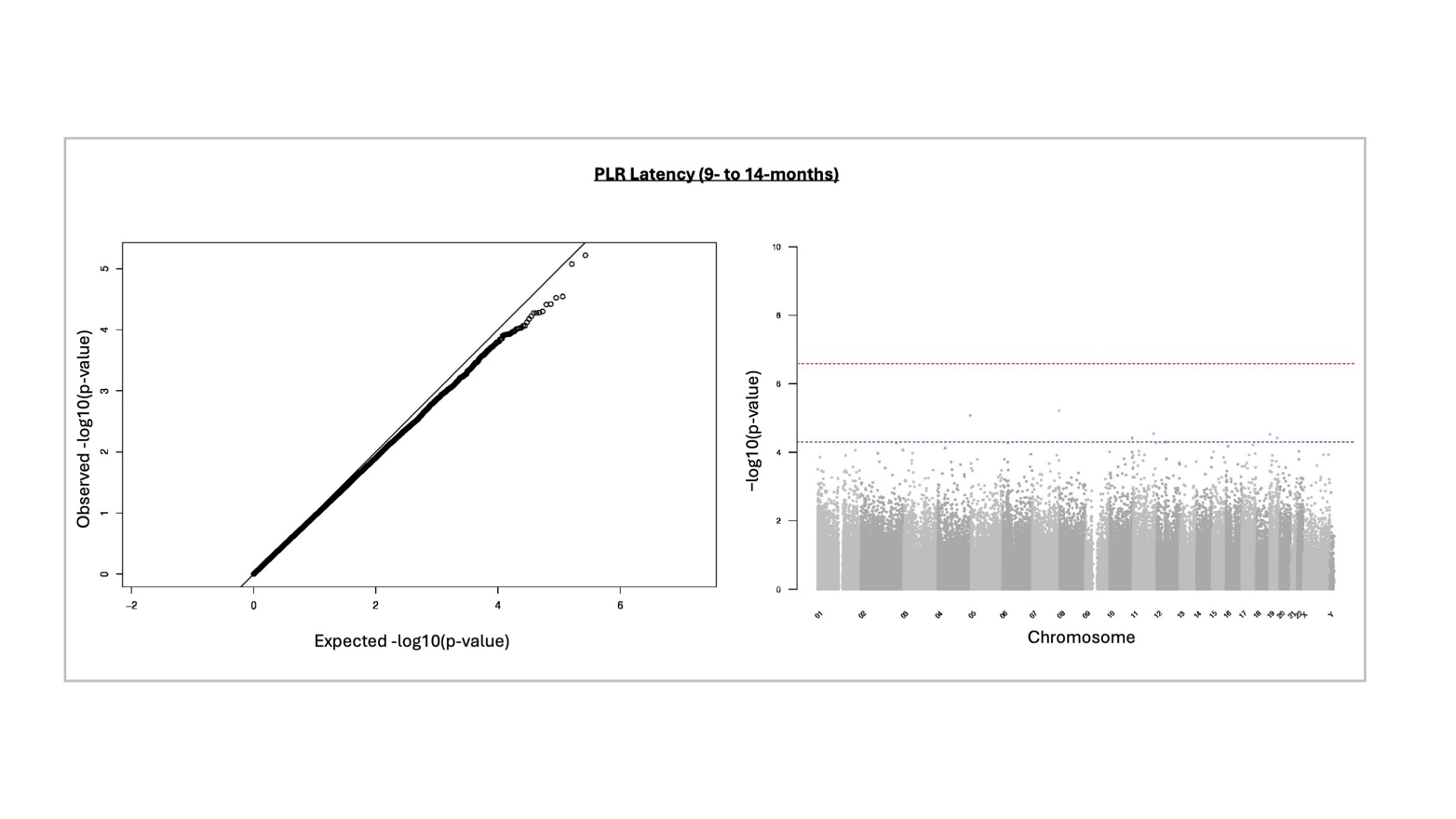

SM 4 Figure 9. Q-Q plot (left) and Manhattan plot (right) of results from 9- to14-month PLR amplitude EWAS

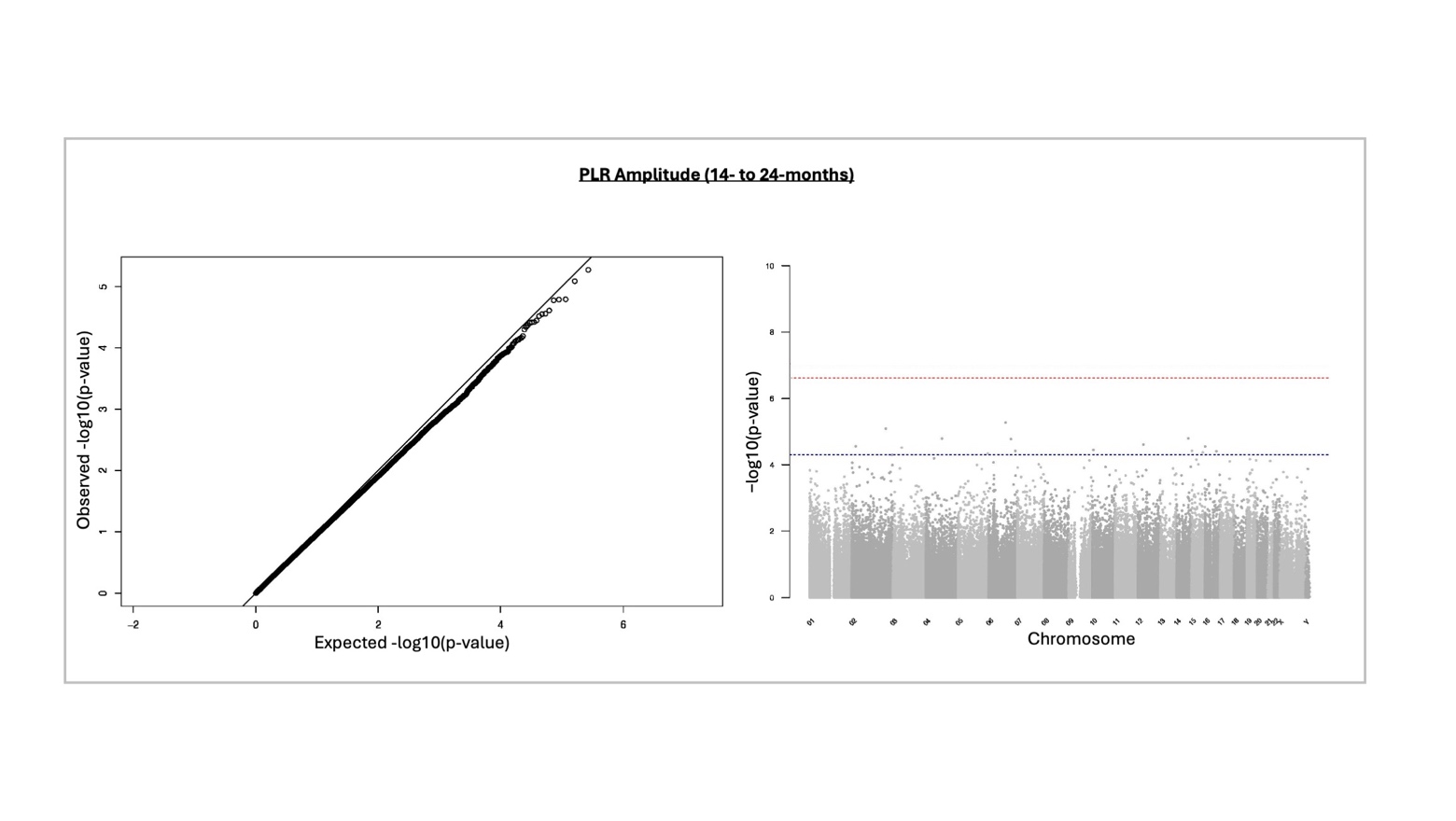

SM 4 Figure 10. Q-Q plot (left) and Manhattan plot (right) of results from 14- to 24-month PLR amplitude EWAS

#### SM 4.2: EWAS probes at discovery p-value threshold (p < 5 × 10^-5^) or above.

See SM 4 Table 1 in accompanying excel document for full list of probes found to be significantly associated in each EWAS discovery p-value threshold *(p* < 5 × 10^-5^) or above.

#### SM 4.3: Significantly associated DMR probes

See SM 4 Table 2 in accompanying excel document for full list of DMR found to be significantly associated.

### SM 5: Downstream exploratory analysis.

#### SM 5.1: Gene Ontology input

See SM 5 Table 1 in accompanying excel document lists genes explored in Gene Ontology analysis for each phenotype.

#### SM 5.2: Gene Ontology results

See SM 5 Table 2 in accompanying excel document lists significant terms found in Gene Ontology analysis for each phenotype.

#### SM 5.3: SFARI gene comparison

See SM 5 Table 3 in accompanying excel document lists probes annotated to genes listed in the SFARI database (with corresponding SFARI gene-score) and associated phenotype.
